# Phosphorus stress induces the synthesis of novel glycolipids in *Pseudomonas aeruginosa* that confer protection against a last-resort antibiotic

**DOI:** 10.1101/2021.04.14.439775

**Authors:** Rebekah A. Jones, Holly Shropshire, Caimeng Zhao, Andrew Murphy, Ian Lidbury, Tao Wei, David J Scanlan, Yin Chen

## Abstract

*Pseudomonas aeruginosa* is a nosocomial pathogen with a prevalence in immunocompromised individuals and is particularly abundant in the lung microbiome of cystic fibrosis patients. A clinically important adaptation for bacterial pathogens during infection is their ability to survive and proliferate under phosphorus (P) limited growth conditions. Here, we demonstrate that *P. aeruginosa* adapts to P-limitation by substituting membrane glycerophospholipids with sugar-containing glycolipids through a lipid renovation pathway involving a phospholipase and two glycosyltransferases. Combining bacterial genetics and multi-omics (proteomics, lipidomics and metatranscriptomic analyses), we show that the surrogate glycolipids monoglycosyldiacylglycerol and glucuronic acid-diacylglycerol are synthesised through the action of a new phospholipase (PA3219) and two glycosyltransferases (PA3218 and PA0842). Comparative genomic analyses revealed that this pathway is strictly conserved in all *P. aeruginosa* strains isolated from a range of clinical and environmental settings and actively expressed in the metatranscriptome of a cystic fibrosis patient. Importantly, this phospholipid-to-glycolipid transition comes with significant ecophysiological consequence in terms of antibiotic sensitivity. Mutants defective in glycolipid synthesis survive poorly when challenged with polymyxin B, a last-resort antibiotic for treating multi-drug resistant *P. aeruginosa*. Thus, we demonstrate an intriguing link between adaptation to environmental stress (nutrient availability) and antibiotic resistance, mediated through membrane lipid renovation that is an important new facet in our understanding of the ecophysiology of this bacterium in the lung microbiome of cystic fibrosis patients.

**Classification:** Integrated genomics and post-genomics approaches in microbial ecology

## Introduction

*P. aeruginosa* is a significant nosocomial pathogen in intensive care units causing pneumonia, surgical wound site infections and sepsis (1-2). It is now recognised as a leading cause of morbidity and mortality in chronically infected cystic fibrosis (CF) patients and immunocompromised individuals due to the surge of carbapenem resistant strains, a key group of first line antibiotics for treating *P. aeruginosa* infections (3). For these drug-resistant *P. aeruginosa* strains, a viable but not ideal treatment option are polymyxins, considered to be last resort antibiotics. Although polymyxins are active against *P. aeruginosa*, their use was originally discontinued due to concerns over toxicity (4). Indeed, *P. aeruginosa* has started to develop mechanisms of resistance to polymyxins due to increase in their use globally. These primarily include modifications to the lipopolysaccharide (LPS) layer of the outer membrane through the addition of 4-amino-4-deoxy-L-arabinose (L-Ara4N) or phosphoethanolamine (pEtN) (5-6). These changes perturb the electrostatic interaction between cationic polymyxins and normally negatively charged LPS.

Glycerophospholipids, such as phosphatidylglycerol (PG) and phosphatidylethanolamine (PE), are the major lipids forming the membrane lipid bilayer in bacteria, archaea and eukaryotes (7-11). They play a fundamental role in the evolution of the cell and it is widely accepted that the last universal common ancestor possessed a phospholipid membrane (12-13). Although it is uncertain why evolution selected glycerophospholipids as the building blocks for maintaining cellular membranes (13), it is known that organisms can alter their membrane lipid composition in response to nutrient stress or environmental changes (7, 14). Previous studies have firmly established the link between nutrient stress, particularly phosphorus (P) availability, and the expression of a variety of virulence factors in *P. aeruginosa* (15-19). However, it is unclear whether adaptation to P limitation in this bacterium causes a change in membrane lipid composition and, if so, whether lipid remodelling comes with unforeseen ecophysiological consequences. Using a synthesis of multi-omics approaches, here we show that *P. aeruginosa* produces surrogate glycolipids to replace phospholipids in response to P limitation. This lipid renovation pathway is strictly conserved in all *P. aeruginosa* strains isolated from a range of clinical settings and actively expressed in the metatranscriptome of a cystic fibrosis patient. Importantly, such a phospholipid-to-glycolipid transition comes with a significant consequence in antibiotic sensitivity, in that glycolipids confer protection when challenged with the antimicrobial peptide polymyxin B. This work highlights more generally how the ‘natural’ physiological adaptation of a pathogen to its *in situ* environment – in this case the P-depleted environment of the human lung – can mediate a physiological response that has profound implications for the survival of the bacteria in the lung microbiome.

## Materials and methods

### Cultivation of *P. aeruginosa* and mutants

*P. aeruginosa* strain PAO1 was obtained from the DSMZ culture collection (Germany) and routinely cultured in lysogeny broth (LB). A defined medium previously outlined for *Pseudomonas* species to control phosphate levels was also used (20). This modified minimal media A comprised: Na-succinate 20 mM, NaCl 200 mg L^-1^, NH_4_Cl 450 mg L^-1^, CaCl_2_ 200 mg L^-1^, KCl 200 mg L^-1^, MgCl_2_ 450 mg L^-1^, with trace metals FeCl_2_ 10 mg L^-1^ and MnCl_2_ 10 mg L^-1^, with 10 mM 4-(2-hydroxyethyl)-1-piperazineethanesulfonic acid (HEPES) buffer used at pH 7. Na_2_HPO_4_ was then added to a final concentration of 50 µM (low P) or 1 mM (high P). An intermediate phosphate source of 400 µM Na_2_HPO_4_ was used for overnight cultures in some experiments to prevent any excess storage of phosphate that could hamper results. All components were filter sterilised using 0.22 µm pore-size filters, and made up using double deionised H_2_O. Mutants were obtained from the *P. aeruginosa* strain PAO1 transposon mutant library at the University of Washington, and confirmed using PCR and subsequent sequencing.

### Alkaline phosphatase assay

Alkaline phosphatase activity was monitored as a measure of phosphate stress. Liquid *P. aeruginosa* culture samples were incubated with 10 mM *para*-nitrophenol phosphate (*p*NPP) to a final concentration of 1 mM. Yellow-*p*NP supernatant was measured in triplicates at 407 nm (BioRad iMark microplate reader). Readings were normalised using both a Tris-only incubation control and further by bacterial density (optical density reading at 600 nm (OD_600_)).

### Over-expression of Agt1 and Agt2 in *E. coli*

*P. aeruginosa* genes PA3218 (*agt1*) and PA0842 (*agt2*) were codon optimised for *E. coli* and chemically synthesised (GenScript) into plasmid pET-28a(+). *E. coli* BLR(DE3) competent cells were thawed for 5 minutes before incubation with 10 ng pET-28a_Agt plasmid, and placed on ice for 5 minutes. Cells were then subjected to heat shock at 42°C for 30 seconds, placed back on ice for 2 minutes. Recovery SOC media was added, with samples incubated at 37°C shaking, for 1 hour. Transformed cells were then plated onto kanamycin-LB agar, grown overnight at 37°C. To harvest cells for lipid extraction, single colonies were picked to grow in small volume LB-Kan to 0.6 OD_600_ before induction with 0.4 mM IPTG overnight at 25°C. 1 mL samples were then pelleted at 10,000 x *g* for 5 minutes. Pellets were stored at -80°C until lipid extraction and subsequent analysis on HPLC-MS.

To purify the Agt1 and Agt2 proteins from recombinant *E. coli*, IPTG was added to a final concentration of 0.5 mM once the cultures reached an OD_600_ of 0.6. After a further 12 h of growth at 30°C, the cells were harvested by centrifugation and resuspended in buffer A containing 50 mM Tris-HCl, pH 7.9, 50 mM NaCl. Cells were disrupted by sonication and 1% (w/v) of triton X-100 was then added which was then incubated for 2.5 hr at 4°C. The cells were then centrifuged at 12,000 × *g* for 20 min, and the soluble fraction was loaded onto a nickel column (GE Healthcare, USA) pre-equilibrated with buffer A. The recombinant Agt1 and Agt2 enzymes were eluted with an elution buffer (20 mM Tris-HCl, pH 7.9, 500 mM NaCl, 300 mM imidazole) and dialyzed overnight into buffer A to remove imidazole. For further purification, the samples were dialyzed overnight into buffer B containing 50 mM Tris-HCl, pH 7.9, 200 mM NaCl, concentrated by ultrafiltration using a 30-kDa membrane (Millipore), and loaded onto a Superdex 200 (16/60) gel filtration column (GE Healthcare, USA), which was pre-equilibrated with buffer B (50 mM Tris-HCl, pH 7.9, 200 mM NaCl). The fraction size was 0.5 ml, and the flowrate was 0.5 ml/min. Purified protein was analysed by SDS-PAGE, and protein concentrations were determined using the Bradford assay.

### Membrane lipid extraction and HPLC-MS analysis

Intact polar membrane lipids were extracted using a modified version of the typically used Folch extraction method (21, 22). Liquid *P. aeruginosa* cultures growing in high and low phosphate modified minimal medium A were sampled after 8 hrs, collecting the equivalent to 0.5 OD_600_ into a 2 mL glass chromacol vial (Thermo Scientific), pelleted at 4°C, 4,000 rpm for 15 minutes. For lipid extraction, a ratio of 500:300:1000 µL of methanol:water:chloroform (all LC-MS grade) was used. The lipid fraction was collected from the lower phase using a glass Pasteur pipette. This chloroform extract was then dried under a stream of nitrogen (Techne sample concentrator) and resuspended in 1 mL 95% (v/v) acetonitrile (HPLC grade): 5% (w/v) ammonium acetate (10 mM, pH 9.2) for analysis. Extracted lipid samples were analysed using an UltiMate 3000 HPLC (Thermo Scientific) system coupled to AmazonSL quadrupole ion trap (Bruker) mass spectrometer (MS), using electrospray ionisation. Hydrophilic interaction chromatography (HILIC) using a BEH amide XP column (Waters) was utilised to separate lipid classes based on their head group (35). The column chamber was maintained at 30°C and the samples passed through at a 150 µL min^-1^ flow rate. The mobile phase of acetonitrile:ammonium acetate (pH 9.2) was used to elute the sample in a 15 minute per sample gradient, from 95% to 28% ammonium acetate. The lipid d17:1/12:0 sphingosylphosphoethanolamine (Sigma-Aldrich, 50 nM) was added to the samples and used as internal standard. Tandem MS (or MS^n^) was used to fragment the intact lipids for identification. The data were analysed using Bruker Compass software package (DataAnalysis and QuantAnalysis).

### Enzyme activity assays

The glycosyltransferase activity of Agt1 and Agt2 was measured using uridine diphosphate (UDP)-glucose or UDP-glucuronic acid and 0.1 mM C16:0/C18:1 diacylglycerol (DAG) as the substrate. 2.0 µM purified enzyme was used in 10 mM Tricine/KOH buffer, pH 8.5 with 2 mM dithiothreitol. The resulting mixture (500 µl) was incubated at 30°C for 60 min with constant shaking at 200 rpm. The lipid products were extracted using the Floch method as described above. The lipid extracts were further analysed by LC-MS for the identification of MGDG/GADG though MS^n^ fragmentation and for the quantification of DAG against standards. The *K*_m_ and *V*_max_ values were calculated using Michaelis-Menten plots with various concentrations of UDP-sugars (0.1 to 1.0 mM) in three replicates.

### Antibiotic sensitivity assays

*P. aeruginosa* cultures were grown to an OD_600_ of 0.6 in high or low phosphate minimal media A (see above). Cultures were then diluted 1:100 in prewarmed minimal media A containing 4 µg mL^-1^ polymyxin B sulfate (Sigma). Samples were incubated at 37°C, 180 rpm, and assayed for survivors at specified time points by serial dilution plating onto LB. *E. coli* cultures containing pET-28a-Agt1 or pET-28a-Agt2 were grown to an OD_600_ of 0.6 in LB broth, and the expression of Agt1 and Agt2 was induced by incubation with 0.4 mM IPTG overnight at 25°C. A negative control of *E. coli* containing the pET-28a vector only was also set up. Overnight cultures were diluted 1:100 in prewarmed LB broth containing 0.4 mM IPTG and 20 µg mL^-1^ polymyxin B sulfate (Sigma). Samples were incubated at 37°C, 180 rpm, and assayed for survivors at specified time points by serial dilution plating onto LB agar + kanamycin 25 µg mL^-1^.

### Comparative proteomic analysis

*P. aeruginosa* PAO1 WT (1 mM phosphate, 50 µM phosphate) and PlcP mutant (50 µM phosphate) cell pellets in three biological replicates were resuspended in LDS (lithium dodecyl sulfate) sample buffer containing 1% β-mercaptoethanol before lysing at 95°C and vortexing. 30 µL of each sample were run on NuPAGE 10% Bis-Tris protein gel (Invitrogen) for a short time before staining with SafeStain (Thermo Fisher) and excising the whole protein band. In-gel proteins were de-stained using 50% (v/v) ethanol, 50 mM ammonium bicarbonate (ABC), before being reduced and alkylated for 5 min at 70°C using 10 mM TCEP (tris(2-carboxyethyl)phosphine) and 40 mM CAA (2-chloroacetamide), respectively. After washing with 50% (v/v) ethanol 50 mM ABC, peptides were lysed overnight using trypsin. Finally, peptides were extracted by sonication in a water bath (10 min at room temperature), concentrated using a Speed-Vac (50 mins) and resuspended in 2.5% acetonitrile 0.05% formic acid. Extracted peptides were analysed by nanoLC-ESI-MS/MS using the Ultimate 3000/Orbitrap Fusion instrumentation (Thermo Scientific). The UniProt proteome for *P. aeruginosa* strain PAO1 was used for peptide analysis. Further data analysis was carried out using MaxQuant and Perseus software; peptides without triplicate measures were filtered out. Comparative proteomics data are presented in supplementary Table S1, S3.

### Phylogenomics and metatranscriptomics analyses

The protein sequences of PA3219, PA3218 and PA0842 were used to search genome sequences of *Pseudomonas* clades in the JGI IMG genome portal (https://img.jgi.doe.gov/). Note that the PA3218 protein is incorrectly annotated in the genome of PAO1. The putative glycosyltransferase located immediately downstream of PA3219 was manually inspected by aligning to the corresponding gene (PA14_22600) in the genome of *P. aeruginosa* PA14. To identify PA3218 in misannotated *P. aeruginosa* genomes, the nucleotide sequence immediately downstream of PA3219 was aligned with PA14_22600. The phylogeny of *Pseudomonas* clades was determined using the nucleotide sequences of six housekeeping genes (*rpoB, rpoD, dnaE, recA, atpD, gyrB*) retrieved from each genome using IQ-Tree with the parameters -m TEST -bb 1000 -alrt 1000. The most suitable model was chosen by the software. Evolutionary distances were inferred using maximum-likelihood analysis. Relationships were visualised using the online platform the Interactive Tree of Life viewer (https://itol.embl.de/). Conserved Pho box sequence was predicted using the MEME server (23).

The metatranscriptomics datasets of sputum samples obtained from a CF patient 7-days (SRX5145606) and 8-days (SRX5145605) before death (24) were retrieved from the short reads archive (SRA) database (https://www.ncbi.nlm.nih.gov/sra). The reads were downloaded using fastq-dump and mapped using the BBMap aligner as described previously (25). Briefly, the SRA reads were mapped to the genome sequence of *P. aeruginosa* PAO1 using a stringent cut-off of minid=0.97. Relative abundance data were compared using RPKM (reads per kilobase of transcript, per million mapped reads). The list of RPKM abundance of individual genes of *P. aeruginosa* PAO1 is shown in **Suppl. Table S4**.

## Results and discussion

### *P. aeruginosa* produces novel glycolipids in response to Pi stress

To determine changes in the membrane lipidome in response to P-stress, the model *P. aeruginosa* strain PAO1 was grown in minimal medium under high (1 mM) or low Pi (50 µM) conditions (**Figure 1a**). The latter condition elicited strong alkaline phosphatase activity, measured through the liberation of *para*-nitrophenol (*p*NP) from *para*-nitrophenol phosphate (*p*NPP) (**Figure 1b**), this being a strong indication that cells were P-stressed. Analysis of membrane lipid profiles using high performance liquid chromatography coupled to mass spectrometry (HPLC-MS) revealed the presence of several new lipids under Pi stress conditions (**Figure 1c**). Thus, during Pi-replete growth (1 mM phosphate), the lipidome is dominated by two glycerophospholipids: phosphatidylglycerol (PG, eluted at 6.8 min) and phosphatidylethanolamine (PE, eluted at 12.2 min). During Pi-stress a lipid species with mass to charge ratio (*m/z*) of 623 and 649 were also found, with MS fragmentation resulting in a 131 *m/z* peak, a diagnostic ion for the amino-acid containing ornithine lipid. This is consistent with previous reports of ornithine lipids in the *P. aeruginosa* membrane in response to Pi stress (26-27).

**Figure 1.**
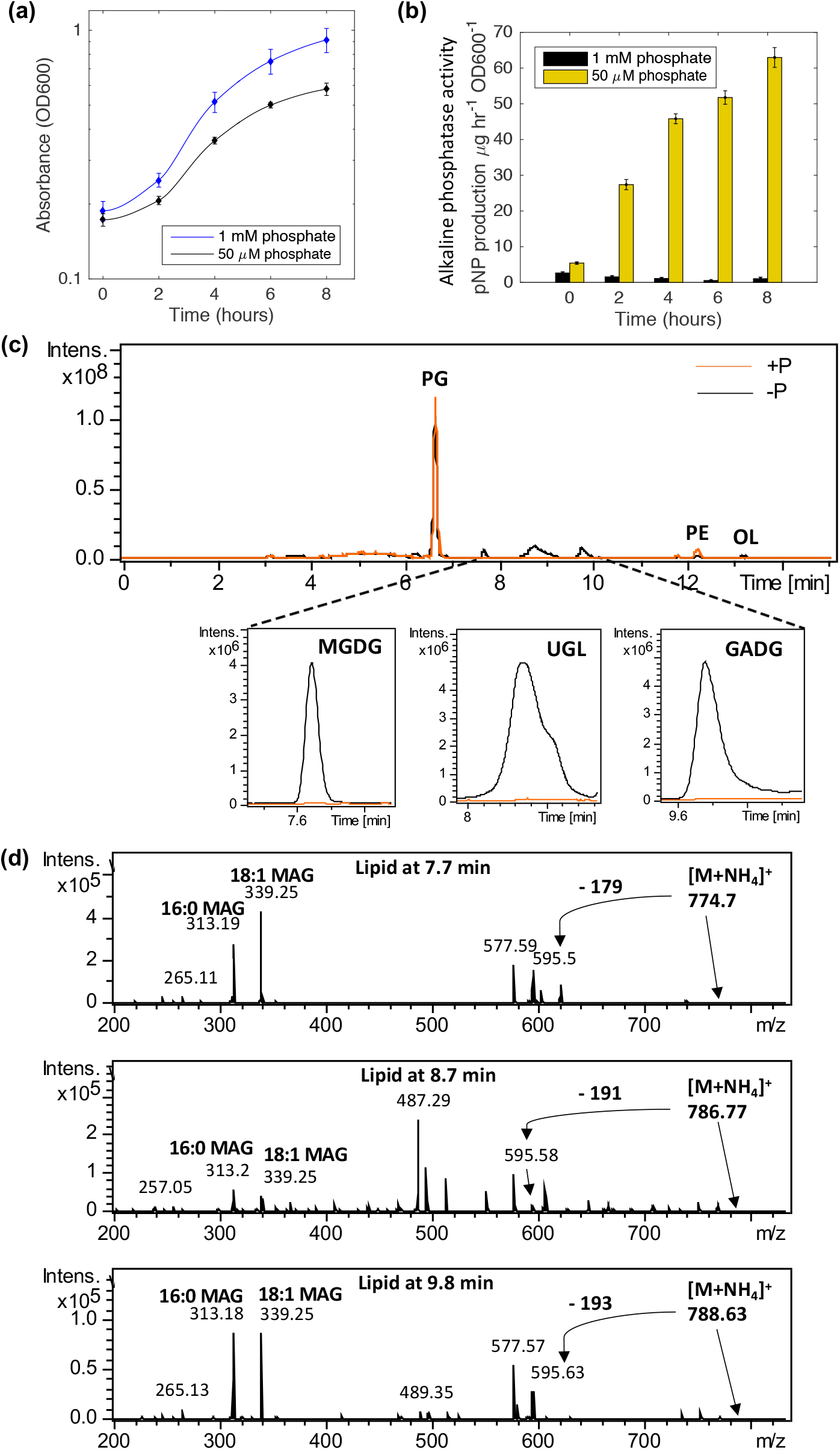
Lipidomics analysis uncovers novel glycolipid formation in *Pseudomonas aeruginosa* strain PAO1 in response to phosphorus limitation. **a)** Growth of strain PAO1 WT in minimal medium A containing 1 mM phosphate (+P, blue) or 50 µM phosphate (-P, black) over 12 hours. Data are the average of 3 independent replicates. **b)** Liberation of *para*-nitrophenol (*p*NP) from *para*-nitrophenol phosphate (*p*NPP) through alkaline phosphatase activity, under P-replete (1 mM, black) and P-deplete (50 µM, yellow) conditions. Error bars represent the standard deviation of three independent replicates. **c)** Representative chromatograms in negative ionisation mode of the *P. aeruginosa* lipidome when grown under phosphorus stress (-P, black) compared to growth under phosphorus sufficient conditions (+P, orange). PG, phosphatidylglycerol; PE, phosphatidylethanolamine, OL, ornithine lipids. Lower panel: extracted ion chromatograms of three new glycolipid species in *P. aeruginosa* which are only produced in P-limitation (black, 1 mM; orange, 50 µM). MGDG: monoglycosyldiacylglycerol, GADG: glucuronic acid-diacylglycerol and UGL: unconfirmed glycolipid. **d)** Mass spectrometry fragmentation spectra of three glycolipid species present under phosphorus stress in *P. aeruginosa*, at retention times of 7.7 (*m/z* 774.7), 8.7 (*m/z* 786.7) and 9.8 (*m/z* 788.6) minutes, respectively. Each spectrum depicts an intact lipid mass with an ammonium (NH_4_^+^) adduct exhibiting neutral loss of a head group, yielding diacylglycerol (DAG) (595 *m/z*). Further fragmentation yields monoacylglycerols (MAG) with C16:0 or C18:1 fatty acyl chains.

Further to ornithine lipids, three unknown lipids eluting at 7.7 min, 8.7 min and 9.8 min, were only present under Pi stress conditions (**Figure 1c**). Using several rounds of MS fragmentation (MS^n^), with a quadrupole ion trap MS, fragmentation patterns characteristic of glycolipids were found for all three peaks. For each peak of interest, the most predominant lipid masses of 774 *m/z*, 786 *m/z* and 788 *m/z* were analysed by MS^n^ in positive ionisation mode (**Figure 1d**). In each case, an initial head group was lost leaving a significant signal of 595 *m/z*, the mass of the glycolipid building block diacylglycerol (DAG). Further fragmentation leads to the loss of either fatty acyl chain from DAG, leaving monoacylglycerols of 313 *m/z* and 339 *m/z*. Two monoacylglycerols with different masses are produced as a result of the original lipid containing 16:0 and 18:1 fatty acids (313 *m/z* and 339 *m/z* monoacylglycerols, respectively). To further elucidate the identity of the peaks, a search for a neutral loss of a polar head group was carried out. Thus, the intact masses of 774 *m/z* and 788 *m/z* in positive ionisation mode leads to the loss of a head group of -179 and -193 *m/z*, which corresponds to a hexose- and a glucuronate-group, respectively (**Figure 1d**), suggesting the occurrence of novel monoglycosyldiacylglycerol (MGDG) and glucuronic acid diacylglycerol (GADG) glycolipids in *P. aeruginosa*. The third glycolipid peak at 8.7 min remains an unknown lipid with intact mass of 767 *m/z* (hereafter designated as a putative unknown glycolipid, UGL). Together, these data confirm the production of new glycolipids in *P. aeruginosa* in response to Pi stress.

### Comparative proteomics uncover the lipid renovation pathway in *P. aeruginosa*

To determine the proteomic response of *P. aeruginosa* to P limitation, and identify the genes involved in glycolipid formation, strain PAO1 was cultivated under high and low Pi conditions for 8 hours and the cellular proteome then analysed. A total of 2844 proteins were detected using nanoLC-ESI-MS/MS instrumentation, 175 of which were found to be differentially regulated by Pi availability when significance was considered at a false detection rate (FDR) of <0.05 (**Figure 2b, Suppl. Table S1**). Of these, 132 proteins were more highly expressed during Pi-deplete conditions, and 43 were more highly expressed during Pi-replete growth conditions. In line with previous transcriptomic studies of *P. aeruginosa* PAO1 (18), major P acquisition mechanisms were more highly expressed under Pi stress conditions. These included the high-affinity Pi-specific transporter PstSCAB, the two-component master regulator of the PHO regulon PhoBR, phosphate-specific porins (OprO, OprP), alkaline phosphatases, phosphonate transporters, extracellular DNA degradation enzymes (EddB and EddA), and a number of key virulence factors (e.g. PlcH, PlcN, PlcB, **Suppl. Table S1**) (28-30).

**Figure 2.**
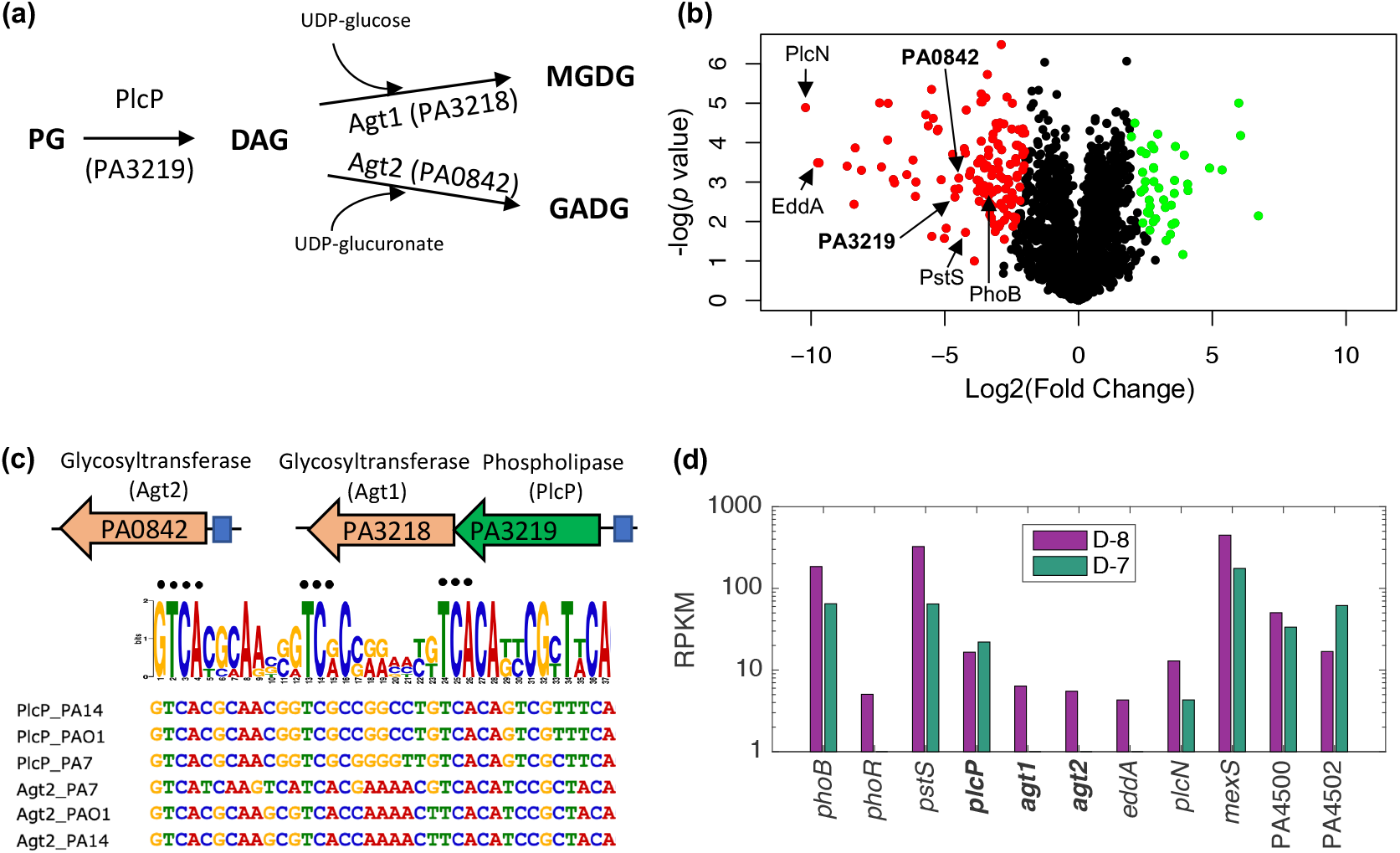
Comparative multi-omic analyses for the identification of the PlcP-Agt pathway responsible for glycolipid formation in *Pseudomonas aeruginosa* strain PAO1. **a)** The proposed pathway for lipid remodelling through the PlcP-Agt pathway. PlcP degrades membrane phospholipids such as PG, to generate diacylglycerol (DAG) intermediates for the formation of MGDG and GADG through the activity of glycosyltransferases, using either UDP-glucose or UDP-glucuronate as the co-substrate (39). **b)** Volcano plot depicting differentially expressed proteins when comparing Pi-replete and Pi-deplete conditions. Significantly upregulated proteins when under Pi stress are shown in red (left), and those that are significantly upregulated when Pi is sufficient are in green (right). Significance was accepted when the false discovery rate (FDR) was < 0.05, and a fold change ≥ 2. **c)** Genomic organisation of predicted lipid remodelling genes in *P. aeruginosa*. Glycosyltransferases (orange) PA3218 (Agt1) and PA0842 (Agt2) are predicted to be involved in glycolipid synthesis. PA3219 is predicted to be PlcP in *P. aeruginosa*. Predicted Pho box sequences in the promoter regions (represented in blue boxes) of each glycosyltransferase operon from *P. aeruginosa* strains representing the PAO1 clade, the PA7 clade and the PA14 clade are shown. The black dots represent residues which are conserved in the Pho box consensus CTGTCATNNNNCTGTCAT (38). **d)** Metatranscriptomic analysis of PlcP-Agt lipid remodelling genes in sputum samples from a cystic fibrosis patient 7-days (D-7) and 8-days (D-8) before death. The relative abundance was expressed as RPKM (reads per kilobase of transcript, per million mapped reads).

Comparative proteomics also identified several genes which are likely important for membrane lipid remodelling (**Figure 2a**) including PA3219 (4.6-fold increase under Pi-deplete conditions, FDR<0.01), encoding a putative phospholipase C protein, and PA0842 (4-fold increase under Pi-deplete conditions, FDR<0.01), encoding a putative glycosyltransferase (**Figure 2b**). PA3219 has 47% protein sequence identity to PlcP from *Phaeobacter* sp. MED193 and 46% identity to PlcP from *Sinorhizobium meliloti* (25-27). In these bacteria, PlcP is essential in the lipid remodelling pathway for the formation of the diacylglycerol (DAG) backbone, representing the essential intermediate for the production of glycolipids (31-32). In *P. aeruginosa* PAO1, PA3219 appears to form an operon with PA3218, a putative glycosyltransferase likely under the control of the PhoBR two component system, as a highly conserved Pho box sequence was recognisable in the promoter region (**Figure 2c**). PA3218 (hereafter referred to as Agt1) has 41% protein sequence identity to the Agt of *Phaeobacter* sp. MED193. PA0842 showed 35% identity to the Agt of *Phaeobacter* sp. MED193 and a Pho box sequence is also found in its promoter region. This corroborates the finding that the PA0842 protein (hereafter referred to as Agt2) was significantly upregulated under Pi-deplete conditions (**Figure 2b**). In summary, comparative proteomic analysis suggests that *P. aeruginosa* PAO1 adopts this PlcP-Agt lipid remodelling pathway for the production of glycolipids in response to Pi-stress (**Figure 2a**).

### The PlcP-Agt mediated lipid renovation pathway is strictly conserved in *P. aeruginosa* and actively transcribed in the metatranscriptomes of a cystic fibrosis patient

To uncover how widespread this predicted PlcP-Agt lipid remodelling pathway is amongst the genus *Pseudomonas*, including *P. aeruginosa* strains, we conducted a thorough comparative genomics analysis of these lipid renovating loci. PlcP-Agt is strictly conserved in all 770 genome-sequenced *P. aeruginosa* strains in the IMG/M database, including all three-previously recognised *P. aeruginosa* lineages (33-34), group 1 represented by strain PAO1, group 2 represented by strain PA14 and group 3 represented by strain PA7 (**Figure 3, Suppl. Table S2**). Indeed, this remodelling pathway is prevalent in many *Pseudomonas* groups, including the plant pathogen *Pseudomonas syringae*. To investigate whether the PlcP-Agt lipid remodelling pathway is involved in host-pathogen interactions, we analysed metatranscriptomic datasets from a cystic fibrosis patient, where *P. aeruginosa* is known to be prevalent in the fatal exacerbation period before patient death (24). To the best of our knowledge, only one study has reported the metatranscriptome of the bacterial community present in CF sputum (24). Indeed, *phoBR* and *pstS* are amongst the most highly expressed genes, confirming previous observations that *P. aeruginosa* is Pi-limited during lung infection (1-2). Importantly, the transcripts of *P. aeruginosa agt1/plcP/agt2* are also highly expressed in CF sputum during the fatal exacerbation period before death (**Figure 2d**). Therefore, our phylogenomic and metatranscriptomic analyses suggest that not only is the PlcP-Agt lipid remodelling pathway strictly conserved and prevalent in *P. aeruginosa*, the genes are also highly expressed during CF patient infection, suggesting a potential role for lipid renovation in host-pathogen interactions.

**Figure 3.**
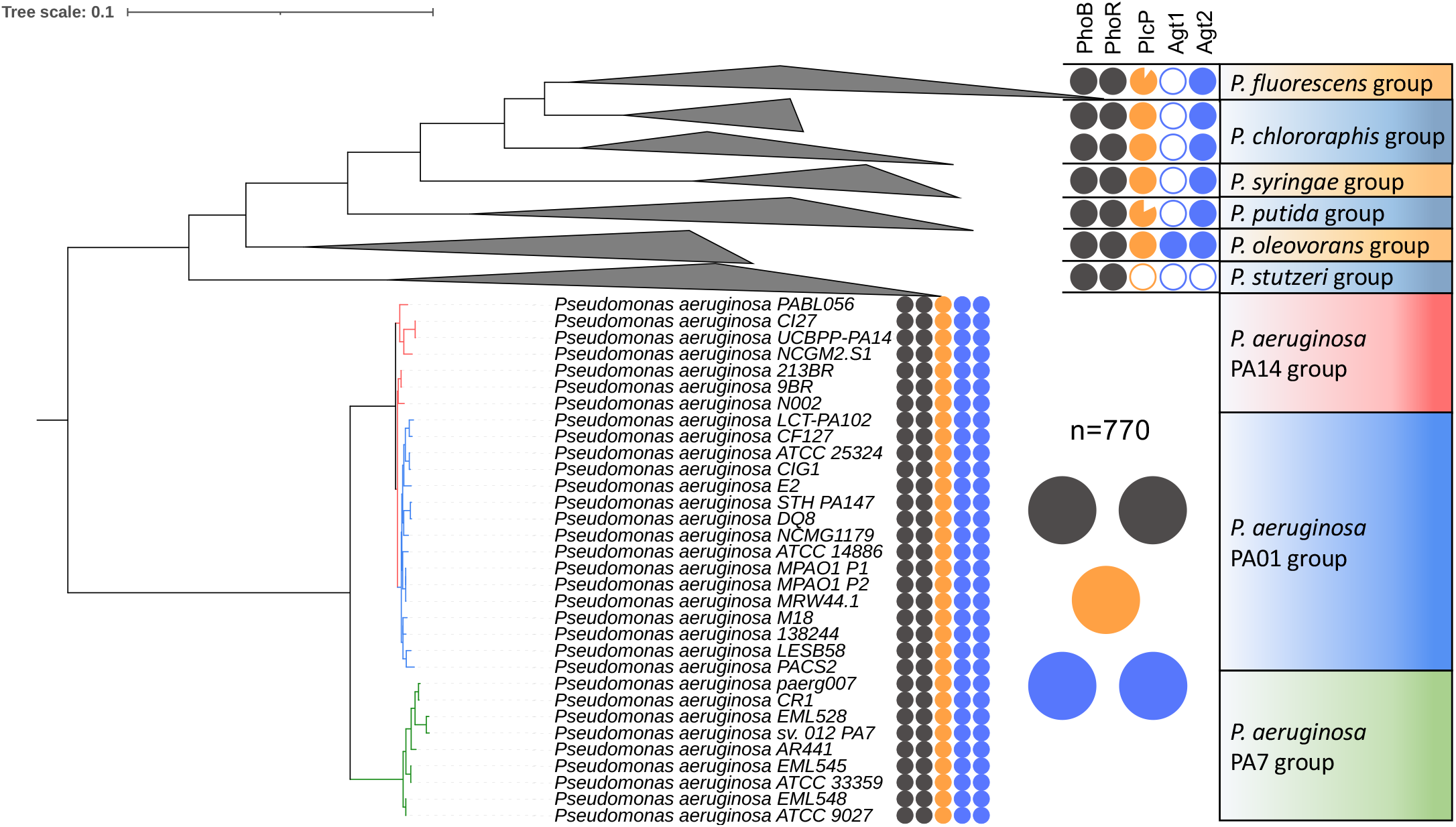
Occurrence of *plcP-agt* genes in major *Pseudomonas* groups. The phylogeny of *Pseudomonas* clades was determined using the nucleotide sequences of six housekeeping genes (*rpoB, rpoD, dnaE, recA, atpD, gyrB*) retrieved from each genome using IQ-Tree (40). The filled colour indicates the presence of the genes in the genomes whereas a blank indicates the absence of the corresponding gene in the genomes. The two-component system PhoBR (black circles) is found in all genomes and the PlcP-Agt1/Agt2 are strictly conserved in all 770 genome-sequenced *P. aeruginosa* strains that form three clades represented by strain PA14, PA01 and PA7 respectively.

### Experimental validation of the lipid renovation pathway for glycolipid formation in *P. aeruginosa*

To validate the function of these two putative glycosyltransferases (Agt1, Agt2) in the formation of glycolipids, we synthesized the codon-optimized genes (PA3218 and PA0842, respectively) for recombinant expression in *Escherichia coli*. The total lipidomes from the recombinant *E. coli* strains were then analysed by HPLC-MS to determine the presence of glycolipids in a gain-of-function assay. Expression of *P. aeruginosa* Agt1 (PA3218) was sufficient for the production of MGDG (eluted at 7.7 min) in *E. coli*, confirmed through MS^n^ fragmentation (**Figure 4a**). No UGL nor GADG was observed in the lipidome of this Agt1-overexpressing *E. coli* strain. Expressing Agt2 (PA0842) from *P. aeruginosa* in *E. coli* was sufficient for the accumulation of the GADG glycolipid (eluted at 9.8 min), also confirmed through the MS^n^ fragmentation pattern (**Figure 4b**). Equally, no UGL nor MGDG was observed in the Agt2-overexpressing *E. coli* strain. Production of these glycolipids was not observed in the same strain of *E. coli* transformed with an empty vector control (pET28a).

**Figure 4.**
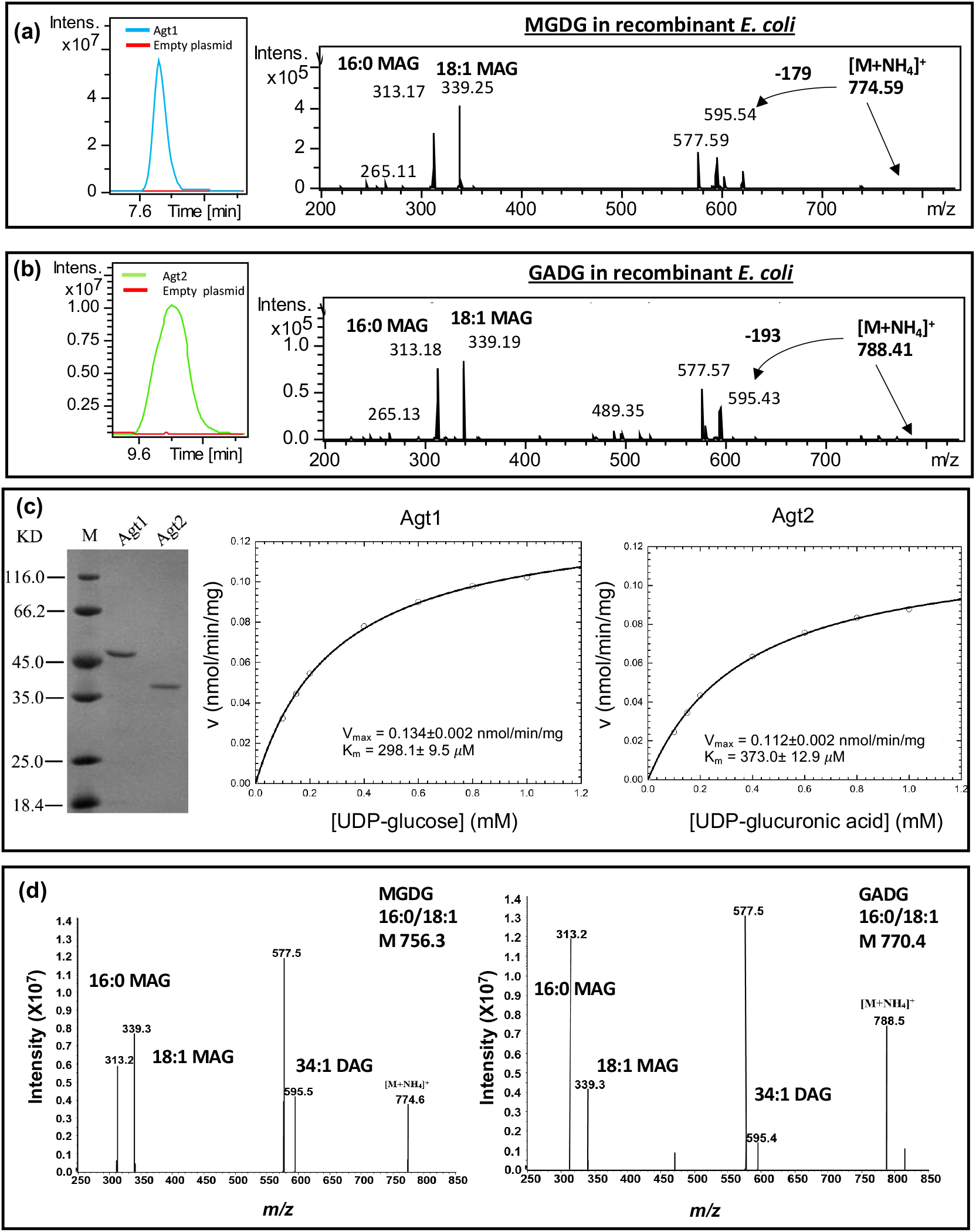
Characterization of glycolipid formation from recombinant Agt1 and Agt2. **a)** Extracted ion chromatogram of the MGDG lipid from recombinant *E. coli* expressing Agt1. An empty vector control is also shown (red line). The identity of MGDG is further validated using mass spectrometry fragmentation showing the neutral loss of 179 corresponding to the loss of glucose and the formation of monoacylglycerols (MAG) with C16:0 or C18:1 (*m/z* 313, 339). **b)** Extracted ion chromatogram of the GADG lipid from recombinant *E. coli* expressing Agt2. An empty vector control is also shown (red line). The identity of GADG is further validated using mass spectrometry fragmentation showing the neutral loss of 193 corresponding to the loss of glucose and the formation of monoacylglycerols (MAG) with C16:0 or C18:1 (*m/z* 313, 339). **c)** Purified Agt1 and Agt2 protein from recombinant *E. coli* (left panel) and Michaelis Menten kinetics of Agt1 towards UDP-glucose (middle panel) and Agt2 towards UDP-glucuronic acid (right panel) as substrate, respectively. **d)** Mass spectrometry identification of MGDG and MADG produced from purified Agt1 and Agt2 using DAG and UDP-glucose and UDP-glucuronic acid as the substrate, respectively.

To confirm the role of Agt1 and Agt2 in the production of MGDG and GADG, we purified Agt1 and Agt2 from recombinant *E. coli* (**Figure 4c**) and carried out enzyme assays using UDP-glucose and UDP-glucuronic acid as the sugar donor and DAG as the acceptor. Agt1 can only accept UDP-glucose as the substrate with an affinity of K_m_=298.1 ± 9.5 *μ*M (**Figure 4c**, middle panel) and produced MGDG as expected (**Figure 4d**, left panel). Similarly, Agt2 can use UDP-glucuronic acid as the substrate (K_m_= 373.0 ± 12.9 *μ*M (**Figure 4c**, right panel), producing GADG lipid (**Figure 4d**, right panel). Interestingly, the purified Agt2 enzyme can also use UDP-glucose to some extent with a K_m_ of 480 *μ*M (data not shown) although the corresponding lipid MGDG was not observed in the lipid extract from the lipidome of the recombinant host *E. coli* (**Figure 4b**).

To further confirm the role of these genes in *P. aeruginosa* glycolipid biosynthesis we analysed the lipidomes of mutants in *ΔplcP, Δagt1* and *Δagt2* in strain PAO1 (**Figure 5a, b**). Differences were analysed by searching for the intact masses of the glycolipids MGDG and GADG: 774 *m/z* and 788 *m/z* in positive ionisation mode with an ammonium adduct, respectively. As expected, under Pi stress MGDG is no longer produced in the *Δagt1* mutant and similarly GADG is no longer produced in the *Δagt2* mutant (**Figure 5a**). In the *ΔplcP* mutant, no MGDG was found and the GADG lipid was significantly reduced but not entirely abolished (**Figure 5b**). The small amount of GADG produced in the *ΔplcP* mutant suggests that an alternative supply of DAG (independent of the degradation of phospholipids by PlcP) is available in this mutant. Nevertheless, lipidome analyses of the *ΔplcP, Δagt1* and *Δagt2* mutants strongly supports the key role of this PlcP-Agt pathway (**Figure 2A**) in lipid renovation in *P. aeruginosa*.

**Figure 5.**
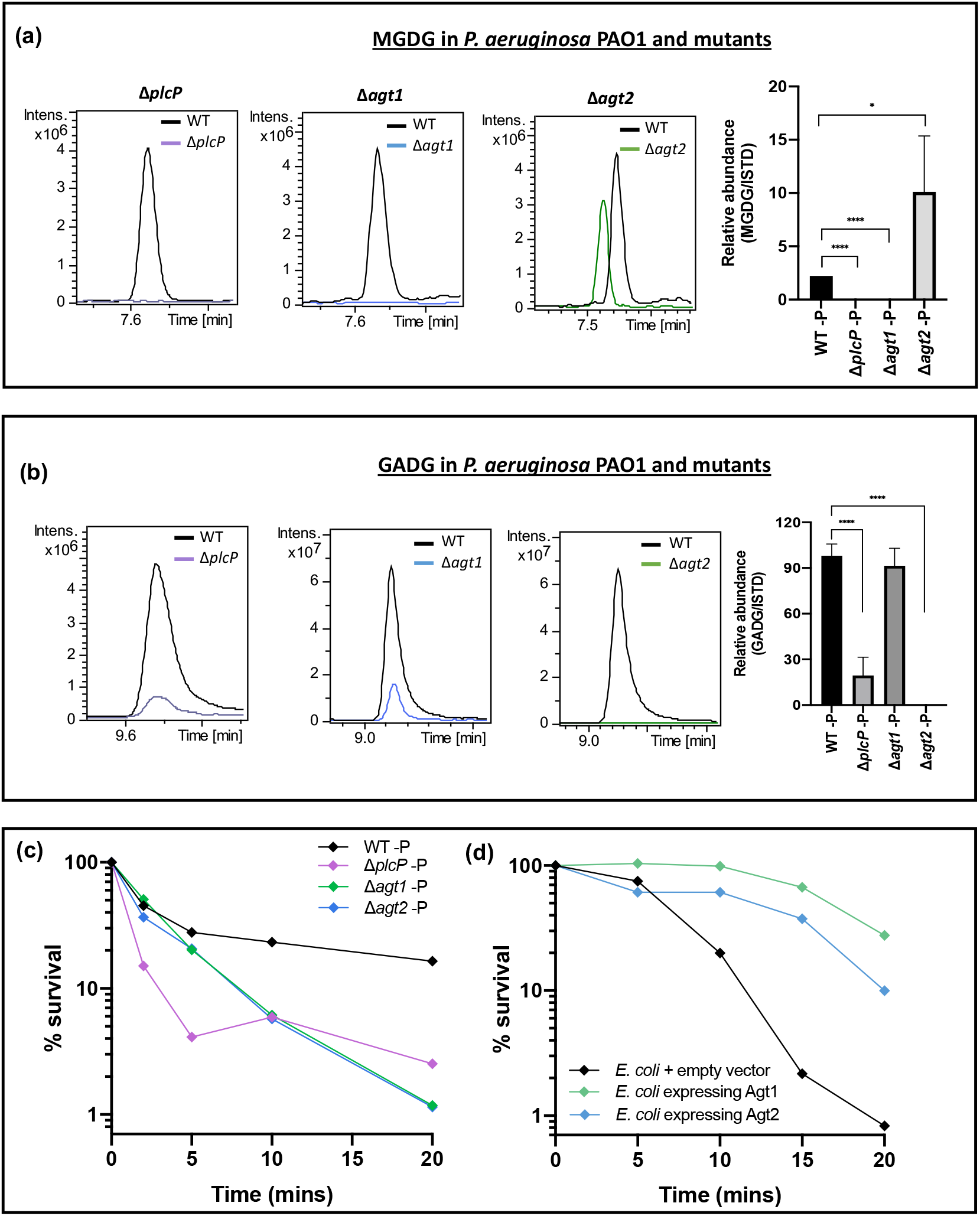
Glycolipid formation in *P. aeruginosa* and mutants under Pi stress, showing a protective role of glycolipids to polymyxinB. **a-b)** Relative abundance of the glycolipid MGDG (**a**) and GADG (**b**) in *P. aeruginosa* mutants *plcP* (purple trace), *agt1* (blue trace) and *agt2* (green, trace) compared to the wild type (WT). Cells were cultivated under low phosphate conditions (50 µM) and a representative extracted ion chromatogram of MGDG/GADG is show between the WT, (black trace) and each mutant. The right most panel shows the abundance of MGDG or GADG calculated relative to an internal lipid standard d17:1/12:0 sphingosylphosphoethanolamine (Sigma-Aldrich) in the wild-type and mutant strains of *P. aeruginosa*. Values are calculated from three biological replicates and the error bars denote standard deviation. MGDG, monoglycosyldiacylglycerol, GADG, glucuronic acid-diacylglycerol. **c)** Survival of glycolipid remodelling mutants Δ*plcP* (purple), Δ*agt1* (green) and Δ*agt2* (blue) when challenged with 4 µg mL^-1^ polymyxin B compared to WT under P stress (black). All experiments were conducted under Pi stress conditions and the results are the average of three biological replicates; error bars denote standard deviation. **d)** Survival of glycolipid producing *Escherichia coli* when challenged with 20 µg mL^-1^ polymyxin B. All experiments were conducted in three replicates and error bars denote standard deviation. Black, *E. coli* containing the empty vector pET28a; green, *E. coli* containing plasmid pET28a-Agt1; blue, *E. coli* containing plasmid pET28a-Agt2.

### The protective role of glycolipids to antibiotic resistance in *Pseudomonas aeruginosa*

The presence of glycolipids in the membrane is likely to have a profound impact on the functioning of the *P. aeruginosa* membrane during Pi stress. For example, PG is an anionic lipid with net negative charges whereas MGDG has a neutral charged sugar group. Although a PG-to-GADG substitution may not necessarily change membrane charge (36), it may affect membrane curvature and the packing density of lipids. Thus, subsequent knock-on effects in membrane function might be expected (10). We therefore set out to investigate whether membrane lipid composition may have an impact on antibiotic resistance in *P. aeruginosa*. As cationic antimicrobial peptides directly interact with bacterial cell membranes, we focused on the impact of lipid remodelling on the killing activity of polymyxin B. We conducted the analyses under P-deplete conditions, since Pi-stress is clinically important, already known to induce the expression of virulence factors (15, 17, 18, 29), and our own analysis confirmed that *P. aeruginosa* is indeed P stressed during CF lung colonisation (**Figure 2d**). Polymyxins represent the drug-of-last resort for effectively treating carbapenem-resistant *P. aeruginosa* infections (3, 37).

To test the sensitivity of the mutants in the PlcP-Agt pathway to polymyxin B, we compared WT and mutants using kill curve analyses as the typically used disk diffusion method does not work efficiently for cationic antimicrobials (38). Indeed, there was a significant decrease in the survival of all three PAO1 glycolipid synthesis mutants (*ΔplcP, Δagt1* and *Δagt2*) compared to the wild type when challenged with polymyxin B, suggesting a protective role of glycolipids in polymyxin B resistance (**Figure 5c**). *P. aeruginosa* is known to enhance its resistance to polymyxins through decoration of its lipopolysaccharide (LPS) layer using either 4-amino-4-deoxy-L-arabinose (L-Ara4N) by *arnB* (5), or the addition of phosphoethanolamine (pEtN) by *eptA* (6). It is thought that these changes perturb the electrostatic interaction between the cationic polymyxin B and the normally negatively charged LPS. To investigate whether these mechanisms play a role in the glycolipid-deficient mutants, we conducted a comparative proteomics analysis of the *ΔplcP* mutant and WT under P deplete conditions, which revealed only a small number of differentially expressed proteins (**Suppl. Table S3**). The majority of these differentially expressed proteins are uncharacterised. However, importantly, LPS modification enzymes previously found to confer antimicrobial peptide resistance, such as ArnB and EptA, were not differentially expressed between the WT and *ΔplcP* mutant. Therefore, the data suggests that these glycolipids are likely a major contributor to increased polymyxin B resistance, which constitutes a new biological mechanism for polymyxin resistance. To this end, we tested the resistance to polymyxin B of recombinant *E. coli* strains overexpressing *P. aeruginosa* Agt1 and Agt2, that produce MGDG and GADG, respectively (**Figure 4a, 4b**). Indeed, in this gain-of-function assay, both Agt1 and Agt2 overexpressing *E. coli* strains had enhanced resistance to polymyxin B compared to the empty vector control (**Figure 5d**), supporting the protective role of these glycolipids to antimicrobial peptides.

To conclude, we present here the discovery of novel glycolipids produced in *P. aeruginosa* during adaption to P stress in the lung microbiome. This lipid renovation pathway is strictly conserved in all *P. aeruginosa* isolates to date and highly expressed in the metatranscriptome of CF patients, suggesting a key role of lipid remodelling in the ecophysiology of this bacterium. Interestingly, lipid remodelling as a response to survive phosphorus stress in turn comes with trade-offs in terms of antibiotic resistance; the glycolipids may protect the bacterium from insult by cationic antimicrobial peptides. It remains to be seen whether the altered susceptibility to polymyxin B is the sole trade-off following lipid remodelling of phospholipids to glycolipids. After all, evolution appears to have selected phospholipids as the dominant lipids in the last universal common ancestor (12).

## Acknowledgements

This work was funded by an MRC Doctoral Training Partnership studentship in Interdisciplinary Biomedical Research (MR/J003964/1) awarded to RAJ and the Royal Society International Exchanges 2017 Cost Share (China) award (IEC\NSFC\170213; grant agreement no. 170213). We thank the Proteomics Research Technology Platform, University of Warwick, UK for their contribution.

## Supplementary Tables

**Suppl. Table S1** Proteomic analysis of differentially expressed proteins in the wild-type *P. aeruginosa* PAO1 in response to different phosphate levels (1 mM versus 50 µM).

**Suppl. Table S2** Protein BLAST identification of locus tags homologous to *agt1* and *agt2* glycolipid synthesis genes in all genome sequenced *P. aeruginosa* strains at the JGI IMG database.

**Suppl. Table S3** Proteomic analysis of wild type *P. aeruginosa* PAO1 versus the Δ*plcP* mutant grown at 50 µM phosphate.

**Suppl. Table S4** Whole genome mapping to *Pseudomonas aeruginosa* PAO1 of a metatranscriptomic dataset (SRX5145605, SRX5145606) from sputum samples taken from a cystic fibrosis patient 7 and 8 days before death. **S4A** (SRX5145605), **S4B** (SRX5145606). RPKM, reads per kilobase of transcript per million mapped reads; FPKM, fragments per kilobase of transcript per million mapped reads.

## References

1. Murray, T. S., Egan, M., & Kazmierczak, B. I. (2007). Pseudomonas aeruginosa chronic colonization in cystic fibrosis patients. Current Opinion in Pediatrics, 19: 83–88.

2. Gaynes, R., Edwards, J. R., & System, N. N. I. S. (2005). Overview of nosocomial infections caused by Gram-negative bacilli. Clinical Infectious Diseases, 41:848– 854.

3. Hawkey PM, Livermore DM (2012). Carbapenem antibiotics for serious infections. BMJ. 344:e3236.

4. Landman, D., Georgescu, C., Martin, D. A., & Quale, J. (2008). Polymyxins revisited. Clinical Microbiology Reviews. 21:449–465.

5. Chung, E. S., Lee, J. Y., Rhee, J. Y., & Ko, K. S. (2017). Colistin resistance in Pseudomonas aeruginosa that is not linked to arnB. Journal of Medical Microbiology, 66:833–841.

6. Nowicki, E. M., O’Brien, J. P., Brodbelt, J. S., & Trent, M. S. (2015). Extracellular zinc induces phosphoethanolamine addition to Pseudomonas aeruginosa lipid A via the ColRS two-component system. Mol. Microbiol. 97:166–178.

7. Parsons JB, Rock CO. (2013) Bacterial lipids: metabolism and membrane homeostasis. Prog Lipid Res. 52:249–276.

8. Zhang, Y.-M., & Rock, C. O. (2008). Membrane lipid homeostasis in bacteria. Nature Rev. Microbiol. 6:222–233.

9. van Meer G, Voelker DR, Feigenson GW. (2008) Membrane lipids: where they are and how they behave. Nat Rev Mol Cell Biol. 9:112–124.

10. Harayama T, Riezman H. (2018) Understanding the diversity of membrane lipid composition. Nat Rev Mol Cell Biol. 19:281–296.

11. May, K.L., & Silhavy T.J. (2017) Making a membrane on the other side of the wall. Biochimica et Biophysica Acta. 1862:1386–1393.

12. Lombard J, López-García P, Moreira D. (2012) The early evolution of lipid membranes and the three domains of life. Nat Rev Microbiol. 10:507–515.

13. Peretó J, López-García P, Moreira D. (2004) Ancestral lipid biosynthesis and early membrane evolution. Trends Biochem Sci. 29:469–477.

14. Sahonero-Canavesi DX, López-Lara IM, Geiger O (2019) Membrane lipid degradation and lipid cycles in microbes. In Aerobic Utilization of Hydrocarbons, Oils, and Lipids, 10.1007/978-3-319-50418-6_38

15. Lamarche MG, Wanner BL, Crépin S, Harel J. (2008) The phosphate regulon and bacterial virulence: a regulatory network connecting phosphate homeostasis and pathogenesis. FEMS Microbiol. Rev. 32:461–473.

16. Long, J., Zaborina, O., Holbrook, C., Zaborin, A., & Alverdy, J. (2008). Depletion of intestinal phosphate after operative injury activates the virulence of P. aeruginosa causing lethal gut-derived sepsis. Surgery, 144:189–197.

17. Francis VI, Stevenson EC, Porter SL. (2017) Two-component systems required for virulence in Pseudomonas aeruginosa. FEMS Microbiol Lett. 364(11). doi:10.1093/femsle/fnx104.

18. Bains, M., Fernández, L., & Hancock, R. E. W. (2012). Phosphate starvation promotes swarming motility and cytotoxicity of Pseudomonas aeruginosa. Applied and Environmental Microbiology, 78:6762–6768.

19. Son, M. S., Matthews, W. J., Kang, Y., Nguyen, D. T., & Hoang, T. T. (2007). In vivo evidence of Pseudomonas aeruginosa nutrient acquisition and pathogenesis in the lungs of cystic fibrosis patients. Infection and Immunity 75:5313–5324.

20. Lidbury, I. D. E. A., Murphy, A. R. J., Scanlan, D. J., Bending, G. D., Jones, A. M. E., Moore, J. D., et al., (2016). Comparative genomic, proteomic and exoproteomic analyses of three Pseudomonas strains reveals novel insights into the phosphorus scavenging capabilities of soil bacteria. Environ. Microbiol. 18:3535–3549.

21. Sebastián, M., Smith, A. F., González, J. M., Fredricks, H. F., Van Mooy, B., Koblížek, M., et al., (2016). Lipid remodelling is a widespread strategy in marine heterotrophic bacteria upon phosphorus deficiency. ISME J. 10:968–978.

22. Smith AF, Rihtman B, Stirrup R, Silvano E, Mausz MA, Scanlan DJ, Chen Y. (2019) Elucidation of glutamine lipid biosynthesis in marine bacteria reveals its importance under phosphorus deplete growth in Rhodobacteraceae. ISME J. 13:39–49.

23. Bailey TL, Elkan C (1994) Fitting a mixture model by expectation maximization to discover motifs in biopolymers. Proceedings of the Second International Conference on Intelligent Systems for Molecular Biology, 28-36, AAAI Press, Menlo Park, Californias.

24. Cobián Güemes AG, Lim YW, Quinn RA, Conrad DJ, Benler S, Maughan H, Edwards R, et al. (2019) Cystic fibrosis rapid response: translating multi-omics data into clinically relevant information. mBio. 16;10(2):e00431–19.

25. Jones, H.J., Krober, E., Stephenson, J., Mausz M.A., Jameson, E., Millard, A., Purdy, K.J., Chen, Y. (2019) A new family of uncultivated bacteria involved in methanogenesis from the ubiquitous osmolyte glycine betaine in coastal saltmarsh sediments. Microbiome. 7, 120. doi.org/10.1186/s40168-019-0732-4

26. Kim, S. K., Park, S. J., Li, X. H., Choi, Y. S., Im, D. S., & Lee, J. H. (2018). Bacterial ornithine lipid, a surrogate membrane lipid under phosphate-limiting conditions, plays important roles in bacterial persistence and interaction with host. Environ. Microbiol. 20:3992–4008.

27. Lewenza, S., Falsafi, R., Bains, M., Rohs, P., Stupak, J., Sprott, G. D., & Hancock, R. E. W. (2011). The olsA gene mediates the synthesis of an ornithine lipid in Pseudomonas aeruginosa during growth under phosphate-limiting conditions, but is not involved in antimicrobial peptide susceptibility. FEMS Microbiol. Lett. 320: 95–102.

28. Kida Y, Shimizu T, Kuwano K. (2011) Cooperation between LepA and PlcH contributes to the in vivo virulence and growth of Pseudomonas aeruginosa in mice. Infect Immun. 79:211–219.

29. Haghi F, Zeighami H, Monazami A, Toutouchi F, Nazaralian S, Naderi G. (2018) Diversity of virulence genes in multidrug resistant Pseudomonas aeruginosa isolated from burn wound infections. Microb Pathog. 115:251–256.

30. Wilton M, Halverson TW, Charron-Mazenod L, Parkins MD, Lewnza S. (2018) Secreted phosphatase and deoxyribonuclease are required by Pseudomonas aeruginosa to defend against neutrophil extracellular traps. Infec Immun 86:e00403–18.

31. Wei, T., Quareshy, M., Zhang, Y. Z., Scanlan, D. J., & Chen, Y. (2018). Manganese is essential for PlcP metallophosphoesterase activity involved in lipid remodeling in abundant marine heterotrophic bacteria. Appl. Environ. Microbiol. 84, e01109–18.

32. Zavaleta-Pastor, M., Sohlenkamp, C., Gao, J.-L., Guan, Z., Zaheer, R., Finan, T. M., et al., (2010). Sinorhizobium meliloti phospholipase C required for lipid remodeling during phosphorus limitation. Proc. Nat. Acad. Sci. USA107:302–307.

33. Freschi L, Jeukens J, Kukavica-Ibrulj I, Boyle B, Dupont MJ, Laroche J, et al., (2015) Clinical utilization of genomics data produced by the international Pseudomonas aeruginosa consortium. Front Microbiol. 29;6:1036. doi:10.3389/fmicb.2015.01036.

34. Ozer EA, Nnah E, Didelot X, Whitaker RJ, Hauser AR. (2019) The population structure of Pseudomonas aeruginosa is characterized by genetic isolation of exoU+ and exoS+ lineages. Genome Biol Evol. 11:1780–1796.

35. Diercks H, Semeniuk A, Gisch N, Moll H, Duda KA, Hölzl G. (2015) Accumulation of novel glycolipids and ornithine lipids in Mesorhizobium loti under phosphate deprivation. J Bacteriol. 197:497–509.

36. Poirel L, Jayol A, Nordmann P. (2017) Polymyxins: antibacterial activity, susceptibility testing, and resistance mechanisms encoded by plasmids or chromosomes. Clin Microbiol Rev. 30:557–596.

37. Ezadi, F., Ardebili A., Mirnead R (2019) Antimicrobial susceptibility testing for polymyxins: challenges, issues, and recommendations. J. Clin. Microbiol. 57, e01390–18.

38. Monds RD, Newell PD, Schwartzman JA, O’Toole GA. (2006) Conservation of the Pho regulon in Pseudomonas fluorescens Pf0-1. Appl Environ Microbiol. 72:1910–1924.

39. Semeniuk A, Sohlenkamp C, Duda K, Hölzl G. (2014) A bifunctional glycosyltransferase from Agrobacterium tumefaciens synthesizes monoglucosyl and glucuronosyl diacylglycerol under phosphate deprivation. J Biol Chem. 289(14):10104–14.

40. Minh BQ, Schmidt HA, Chernomor O, Schrempf D, Woodhams MD, von Haeseler A, et al. (2020) IQ-TREE 2: New models and efficient methods for phylogenetic inference in the genomic era. Mol Biol Evol. 37:1530–1534.

